# Nuclear Morphology Optimized Deep Hybrid Learning (NUMODRIL): A novel architecture for accurate diagnosis/prognosis of Ovarian Cancer

**DOI:** 10.1101/2020.11.23.393660

**Authors:** Duhita Sengupta, Sk Nishan Ali, Aditya Bhattacharya, Joy Mustafi, Asima Mukhopadhyay, Kaushik Sengupta

## Abstract

Nuclear morphological features are potent determining factors for clinical diagnostic approaches adopted by pathologists to analyse the malignant potential of cancer cells. Considering the structural alteration of nucleus in cancer cells, various groups have developed machine learning techniques based on variation in nuclear morphometric information like nuclear shape, size, nucleus-cytoplasm ratio and various non-parametric methods like deep learning have also been tested for analysing immunohistochemistry images of tissue samples for diagnosing various cancers. Our aim is to study the morphometric distribution of nuclear lamin proteins as a specific parameter in ovarian cancer tissues. Besides being the principal mechanical component of the nucleus, lamins also present a platform for binding of proteins and chromatin thereby serving a wide range of nuclear functions like maintenance of genome stability, chromatin regulation. Altered expression of lamins in different subtypes of cancer is now evident from data across the world. It has already been elucidated that in ovarian cancer, extent of alteration in nuclear shape and morphology can determine degree of genetic changes and thus can be utilized to predict the outcome of low to high form of serous carcinoma. In this work, we have performed exhaustive imaging of ovarian cancer versus normal tissue and introduced a novel Deep Hybrid Learning approach on the basis of the distribution of lamin proteins. Although developed with ovarian cancer datasets in view, this architecture would be of immense importance in accurate and fast diagnosis and prognosis of all types of cancer associated with lamin induced morphological changes and would perform across small/medium to large datasets with equal efficiency.

**Significance Statement:** We have developed a novel Deep Hybrid Learning approach based on nuclear morphology to classify normal and ovarian cancer tissues with highest possible accuracy and speed. Ovarian cancer cells can be easily distinguished from their enlarged nuclear morphology as is evident from lamin A & B distribution pattern. This is the first report to invoke specific nuclear markers like lamin A & B instead of classical haematoxylin-eosin staining in an effort to build parametric datasets. Our approach has been shown to outperform the existing deep learning techniques in training and validation of datasets over a wide range. Therefore this method could be used as a robust model to predict malignant transformations of benign nuclei and thus be implemented in the diagnosis and prognosis of ovarian cancer in future. Most importantly, this method can be perceived as a generalized approach in the diagnosis for all types of cancer.

## Introduction

The nuclear envelop is a double membrane structure encapsulating the lamina which is a thick meshwork of A & B-type lamin proteins. Nuclear lamins are type V intermediate filament proteins which impart proper size and shape to the nucleus thus conferring mechanical stability (*1*). Lamins also provide a scaffold for binding of several proteins and chromatin. Lamins are associated with a wide range of nuclear functions like nuclear stability (*2*), genome organisation (*3, 4*), protein interaction (*5, 6*), DNA damage repair (*7, 8*), intracellular signalling(*9-11*) and it has got vital roles in replication (*12, 13*), transcription(*14-16*), and splicing(*17, 18*) as well. The mammalian genome encodes three genes for lamin proteins namely, LMNA, LMNB1 and LMNB2. LMNA is alternatively spliced into two major isoforms lamin A and lamin C and two minor isoforms, lamin C2 and AΔ10. LMNB1 codes for lamin B1 and LMNB2 codes for lamin B2 and lamin B3, the expression of human lamin B3 is restricted to the male germ line (*19*). All these three types of lamin A, B and C form discrete self-interacting, independent meshwork but interconnected lattice structures with different physical properties(*20-23*).In mammals, B type lamins are ubiquitously expressed in all tissue types but lamin A and C are developmentally regulated and expressed in differentiated cells (*24*). Lamin A is not detectable in the embryonic stage but its expression sets in when the cells begin to proliferate and divide (*24*). Studies in monoblastoid cells have shown that induction of differentiation is associated with enhanced expression of lamin A/C along with lateral observations like reduced cell proliferation and stronger substratum contact. However, upon withdrawal of the inducer, proliferation kinetics was restored indicating retro-differentiation which was then accompanied by decreased levels of lamin A/C (*25*). These studies lead to the hypothesis that lamin A/C might stabilize differentiation and thus its absence was a major driving force for rapid division in various tumor cells (*26-28*). But in many tumor types, lamin A/C levels are found to be increased which in turn is associated with their aggressive metastatic potential, which cannot be explained by retro-differentiation (*27*). Rather in these cases increased level of lamin A/C might play vital roles in tumour progression by helping the cells overcome the mechanical stress and resist DNA damage induced cell cycle arrest as it may also help in recruitment of DNA damage repair proteins. In either cases, change in expression levels of lamin A/C largely remains a reliable prognostic marker for different tumor types and stages(*29*). Change in the karyoplasmic ratio is a common phenomenon in various types of cancer. But the correlation of this change with tumorigenesis is still not clearly elucidated. There has been a report which attributes molecular crowding events to higher degree of genetic rearrangements leading to nuclear enlargement(*30*). Being the major architectural protein of animal cell nuclei, lamins must be playing a vital role in this alteration of size(*31*). Loss of lamin or their mutations leading to deformed nuclear morphology has been widely studied by different groups(*1, 32, 33*). Interestingly, the differential expression pattern of lamins in different cancers has also been documented from researches across the world (*34*). Recent findings have demonstrated methods to study nuclear morphologies in the light of fluorescent imaging and deep learning (*35*). Radhakrishnan et al have implied this idea in the breast cancer model and proposed a convolutional neural network pipeline which discriminated between normal and cancer cells with high accuracy (*35*). Several other groups have implied machine learning algorithms to classify stages and types of ovarian and breast carcinoma based on broad cytological differences between tissue specimens stained with Haematoxylin-Eosin stain (*36-39*). But these data were essentially non-parametric or in other words, not based on specific annotations of prognostic markers.

In this work, our aim is to study the morphometric distribution of nuclear lamins in ovarian cancer tissues from patients. Being a resident of the nuclear periphery, lamin A and B, when visualized and imaged under confocal microscope in different ovarian cancer tissue samples, can give us information about the shape and size of the nucleus. In the present study on ovarian cancer model, we have introduced lamin A and B as specific markers for classification and accordingly designed an advanced deep hybrid learning (DHL) architecture which helped us extract information from the tissue types and classify them accordingly by deep hybrid learning algorithm which in turn can aid in pathological inspections proposing the ratio of lamin A and B to be clinically significant for diagnosing ovarian cancer. Consequently, this could be implemented to predict the onset and progression of ovarian cancer respectively. Systematic analyses of patient tissue samples in the form of tissue microarray (TMA) were accomplished by confocal imaging. This dataset of images were used as input for developing a fast and accurate deep hybrid learning method.

## Results

### Distinct nuclear morphology of ovarian cancer tissues

Formalin fixed paraffin embedded normal and diseased (ovarian cancer) tissues were obtained from Tata Medical Centre following the ethical guidelines. Tissues were stained with lamin A, and lamin B following proper antigen retrieval technique and imaged under confocal microscope. 15 fields of view, each containing approximately 100 nuclei were captured from each of the subsets (lamin A stained Normal and Ovarian Cancer tissues, lamin B stained Normal and Ovarian Cancer tissues) under similar acquisition parameters. One representative field from each of the tissue sets has been shown in **Figure 1**. A visibly prominent enlargement of the cancer nuclei was observed with respect to the normal nuclei in both lamin A and lamin B stained tissues. Two tissue microarrays each containing 40 samples were obtained from Tata Medical Centre for which we were blinded. The arrays were stained for lamin A and lamin B following the same procedure and consequently the images acquired from the TMA slides were used for validation of best working model for this problem. 40 samples in a tissue microarray slide stained with lamin A antibody has been shown in **Figure 2**.

**Figure 1:**
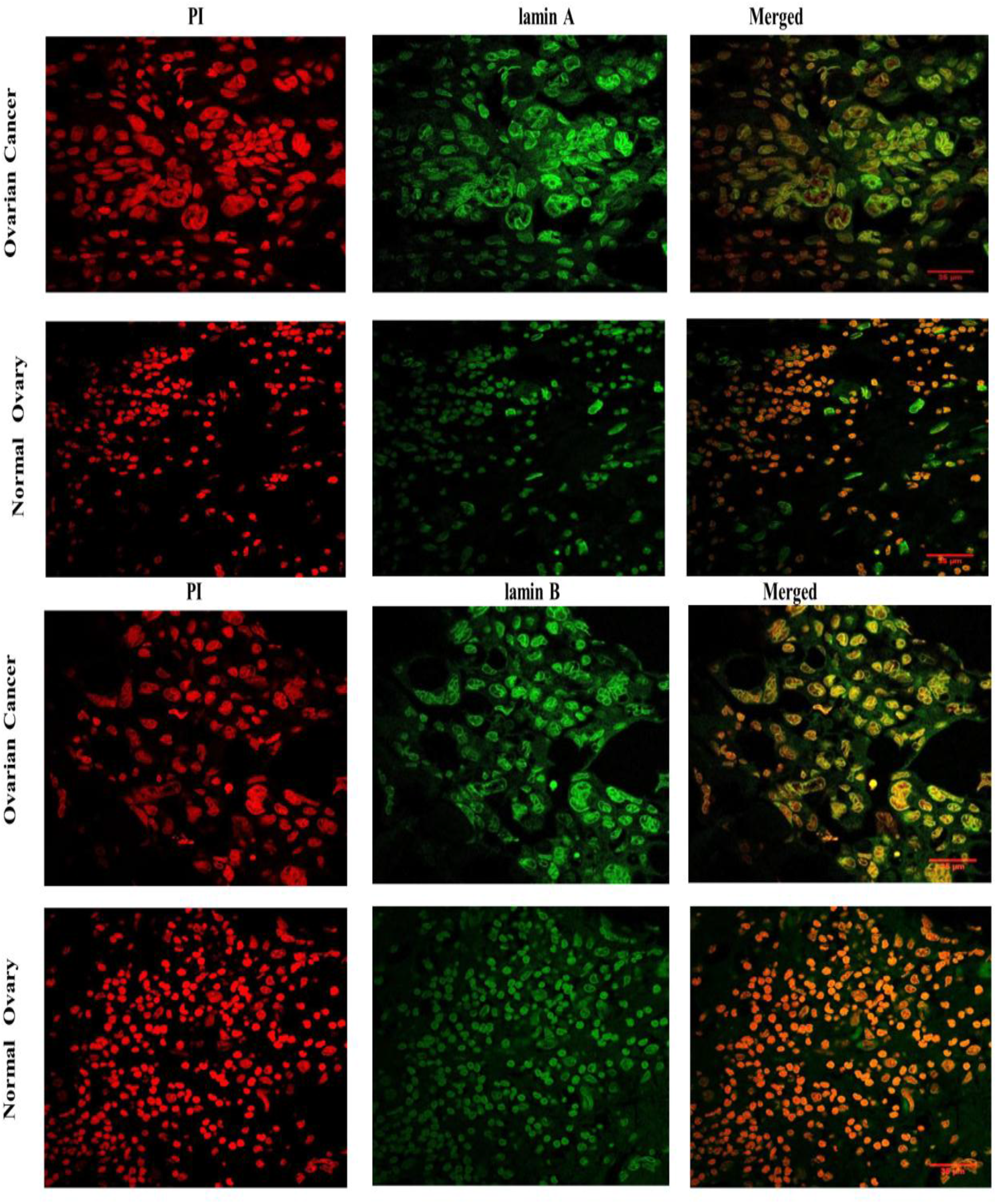
Representative Fields of confocal micrographs showing distribution of lamin A and lamin B in tissues from Ovarian cancer and Normal Ovary . Images of Ovarian Cancer and Normal ovarian tissue nuclei have been marked in their respective columns. Propidium Iodide staining of the nuclei is shown in the first panel containing red channel images. Lamin A and lamin B distributions respectively have been shown in the second panel of green channel images. Merged images of both the channels have been shown in the third panel. Magnification: 63X. Scale Bar: 35 µm

**Figure 2:**
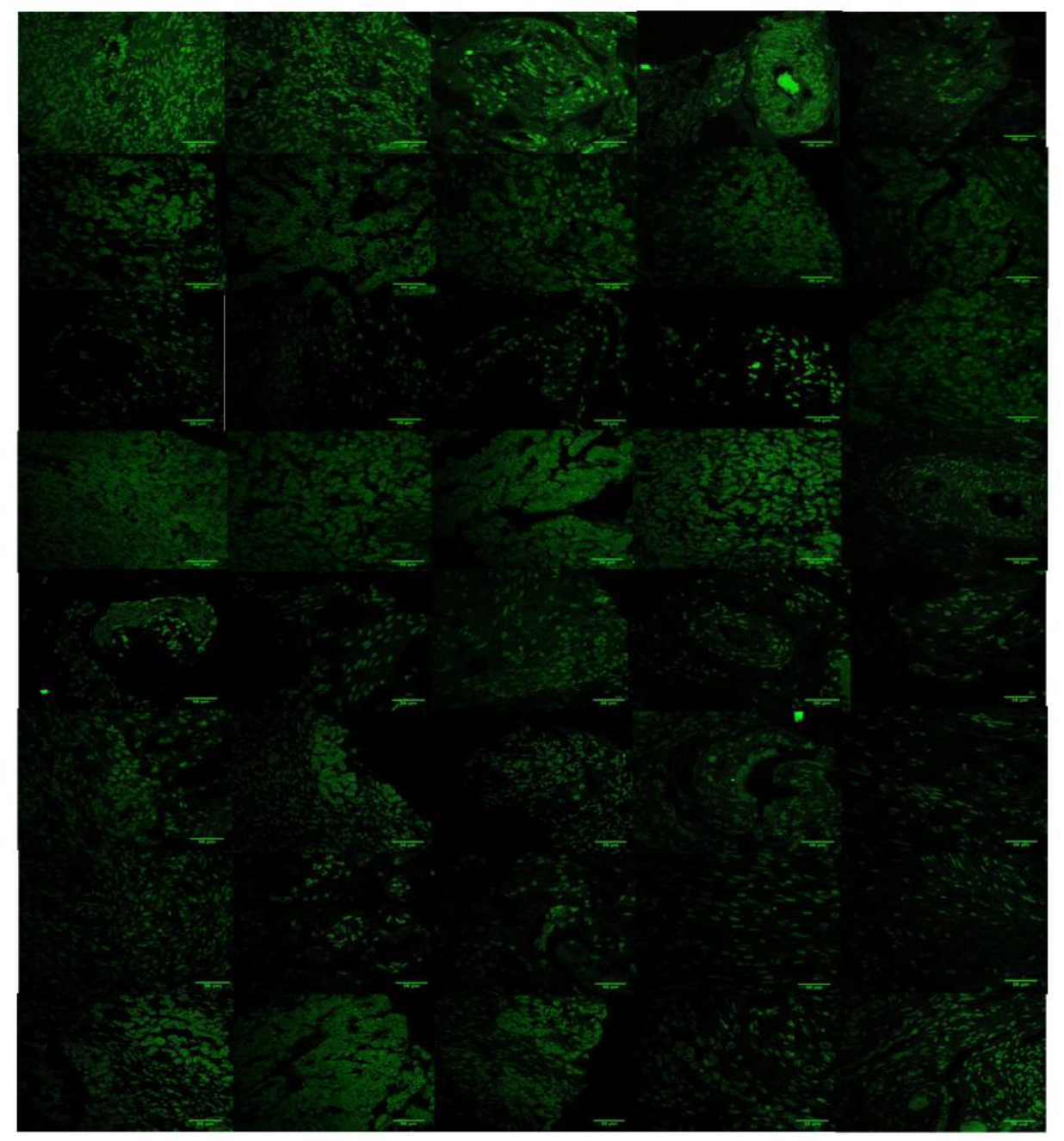
40 samples in a tissue microarray slide stained with lamin. **A**. Each field has been captured under Magnification: 63x. Scale Bar: 35 µm

### Data Augmentation by SMOTE and Analysis of data points in the sample space

We started our experiment with 262 fields each containing about 150 nuclei of ovarian cancer tissues (majority class) and 52 fields each containing about 110 nuclei of normal ovarian epithelial tissue (minority class). Considering the dataset used, the distribution between the majority and the minority class was not equal and the majority class comprised of almost 84% of the dataset, and hence the dataset used was an imbalanced one. Using an imbalanced dataset for building a Deep Learning classifier would add bias to the majority class and unless the dataset is synthetically matched, the classifier would have a strong tendency to predict the majority class for unknown samples. Therefore, we have applied Synthetic Minority Oversampling Technique (SMOTE)(*40*), which uses vector interpolation with high dimensional data to generate synthetic samples of the minority class. Implications of different properties of SMOTE over high dimensional data has been shown in **Supplementary Table 1**.

**Table 1:**
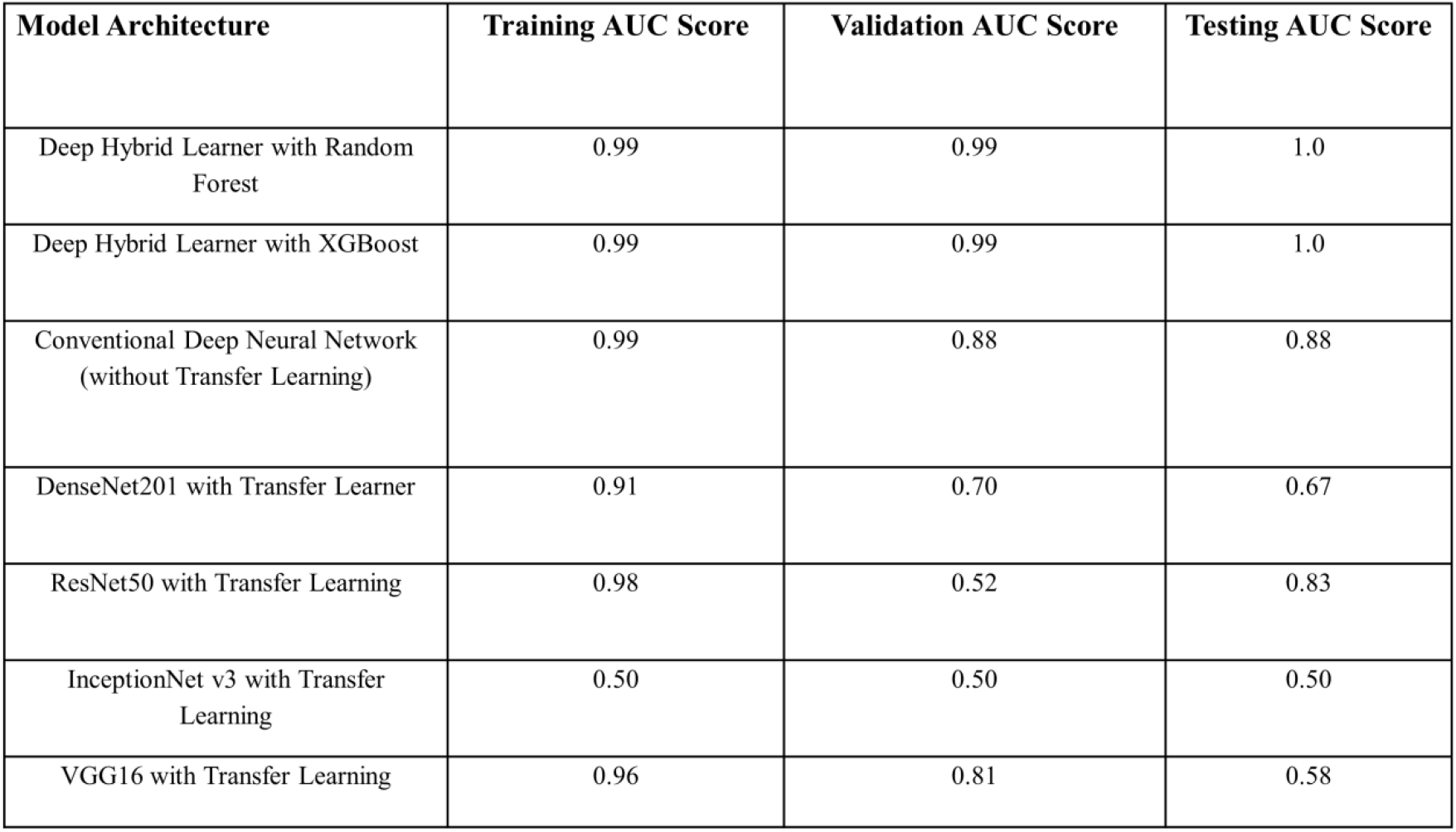
Comparison of model evaluation matrices

| Model Architecture | Training AUC Score | Validation AUC Score | Testing AUC Score |
| --- | --- | --- | --- |
| Deep Hybrid Learner with Random Forest | 0.99 | 0.99 | 1.0 |
| Deep Hybrid Learner with XGBoost | 0.99 | 0.99 | 1.0 |
| Conventional Deep Neural Network (without Transfer Learning) | 0.99 | 0.88 | 0.88 |
| DenseNet201 with Transfer Learner | 0.91 | 0.70 | 0.67 |
| ResNet50 with Transfer Learning | 0.98 | 0.52 | 0.83 |
| InceptionNet v3 with Transfer Learning | 0.50 | 0.50 | 0.50 |
| VGG16 with Transfer Learning | 0.96 | 0.81 | 0.58 |

### Image Pre-processing

The pre-processing algorithm consists of two parts – applying a segmentation mask for cancer cells based on the key visual properties like area, perimeter, circularity, eccentricity, foci distance, loop length, maximum curvature and normalised curvature of the nuclei followed by image sharpening techniques. In the first part, based on the Image Hue Saturation Value (HSV) and using a sensitivity factor, the segmentation mask was created, which was subsequently made prominent by the application of morphological closing operation with an elliptical kernel .Elliptical kernel was used to adapt to the shape of the cells and capture the maximum possible relevant information. Rectangular kernel was previously tested and it got no more than 93% accuracy. In the second phase, using Gaussian Blur and adding weights to the blurred image, we ensured uniform sharpening of the pre-processed images with the segmentation masks and converted the pre-processed images into grey-scaled form so that the key visual features are made more prominent and easier for the Deep Learning algorithm to unravel features. Information from the background was completely removed to emphasize over the morphological properties of the nuclei. (**Figure 3**)

**Figure 3:**
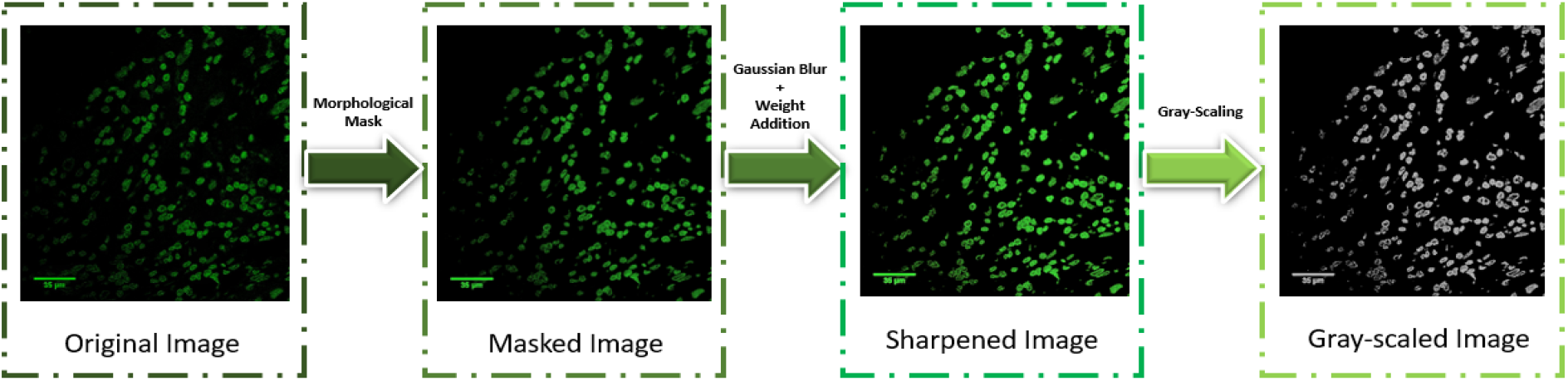
Sequence of operations followed for pre-processing of the image dataset: In the first part, segmentation mask was applied based on the morphometric properties followed by image sharpening to have the key features more visually prominent. In the second part, the sharpened image, which was obtained using Gaussian Blurring and weight addition was transformed to its gray scale version to improve the computational time of the model.

### Morphometric comparisons between images before and after pre-processing

15 fields of view, each containing approximately 100 nuclei were captured from each lamin A and lamin B stained tissue samples under similar acquisition parameters.The minor and major axes of each nucleus were measured manually using ImageJ (ImageJ bundled with 64-bit Java 1.8.0_112). Careful investigation revealed that the hallmark of the diseased tissues was characterized by prominent nuclear enlargement as reported earlier(*41-43*). We quantified these changes by considering every nucleus an ellipse; eight parameters (Area, Perimeter, Eccentricity, Circularity, Foci Distance, Loop Length, Maximum Curvature and Normalized Maximum Curvature) were measured for each of the nuclei using the formulae mentioned earlier (*44*). With these sets of images, a gross morphometric analysis was performed based on the distribution of lamin A and lamin B proteins in the nucleus. Later on, following pre-processing, all the eight parameters were reanalysed from the lamin A and lamin B stained nuclei on randomly selected cancer and normal pre-processed tissue images. Histograms were generated for each of the parameters using ROOT data analysis framework (Version 6, Release 6.08/06-2017-03-02), where the X axis denotes the normalized number of nuclei with respect to the total number of nuclei calculated corresponding to the defined parameter and Y axis denotes the measure of the parameter. It is evident from the plots (**Figure 4**), that the perimeter of most of the cancer nuclei from the total population are showing an increase of 55-62% compared to most of the normal nuclei for both lamin A and lamin B stained tissues in the images before pre-processing and the observation is similar in the pre-processed counterparts as well **(Figure 4 A1, A2)**. Same has been observed while measuring the area, where the area of most of the cancer nuclei is more than twice the area of most of the normal nuclei in the population in both raw and pre-processed images of cancer and normal tissue **(Figure 4 B1, B2)**. Both the observations indicate an increase in size of the cancerous nuclei and this feature is unaltered post pre-processing. However, in the cancer nuclei, around 3% and 12% shift from the normal were observed in the circularity and eccentricity values respectively which is not that significant denoting no prominent change in the shape **(Supplementary figure 1 A1,A2, B1, B2)**. Eccentricity is a focal length (Distance from the centre to one focus) and semi major axis dependent variable. Still, to further validate, foci distance (2*Focal length) was also measured where the shift associated with eccentricity was supposed to get doubled according to the formulae and we could find a small increase in the Foci distance of the cancer nuclei in comparison to the normal nuclei, which denotes an increase in the distance between the foci thereby approaching an elliptical nature **(Supplementary figure 1 C1, C2)**. Another common parameter in ellipse geometry is loop length, which is a focal length dependent variable, hence a rise was evident in the loop length of cancer nuclei denoting an increase in size once again **(Supplementary figure 1 D1,D2)**. Next, to study the change in the surface architecture, maximum curvature and normalised curvature were measured; but no significant shift was observed to deduce a conclusion **(Supplementary figure 1 E1, E2, F1, F2**). Observations were consistent in the pre-processed images as well. As we all know, that tumor microenvironment harbours a heterogeneous cell population including cells at different stages of malignancy and some normal cells too, so the analysis spanned a large range of parametric measures to accommodate all the nuclei in the population. Some shifts were visibly clear and prominent but some were not. So, the change may not be specified with distinct values.

**Figure 4:**
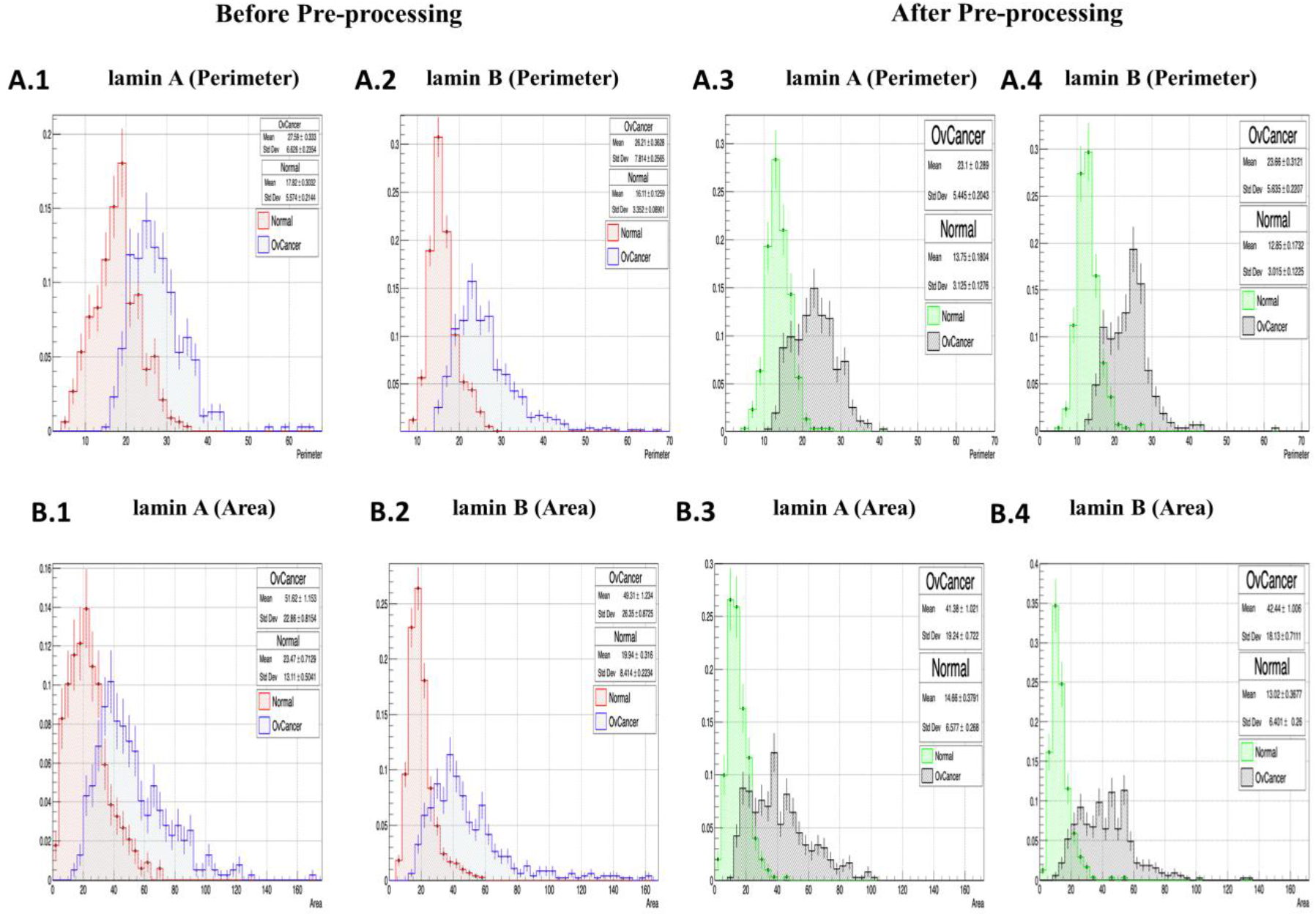
Histograms showing distributions of the normal and Ovarian Cancer nuclei based on different morphometric parameters obtained from lamin A and B stained tissue sample images before and after pre-processing. X axis denotes the normalised number of nuclei with respect to the total number of nuclei calculated. Y axis denotes the measure of the parameter. **A. 1**.Comparative distribution of the number of normal (Mean±Std error of mean:17.82± 0.3032) (Std Dev:5.574±0.2144) and ovarian cancer (Mean±Std error of mean:27.59± 0.333) (Std Dev:6.626±0.2354) nuclei based on Perimeter values acquired from lamin A stained tissue images before pre-processing. **A. 2**. Comparative distribution of the number of normal (Mean±Std error of mean:16.11± 0.1259) (Std Dev:3.352±0.08) and ovarian cancer (Mean±Std error of mean:26.21± 0.3628) (Std Dev:7.814±0.2565) nuclei based on Perimeter values acquired from lamin B stained tissue images before pre-processing. **A. 3**.Comparative distribution of the number of normal (Mean±Std error of mean:13.75± 0.1804) (Std Dev:3.125±0.1276) and ovarian cancer (Mean±Std error of mean:23.1± 0.289) (Std Dev:5.445±0.2043) nuclei based on Perimeter values acquired from lamin A stained tissue images after pre-processing. **A. 4**. Comparative distribution of the number of normal (Mean±Std error of mean:12.85± 0.1732) (Std Dev:3.015±0.1225) and ovarian cancer (Mean±Std error of mean:23.66± 0.3121) (Std Dev:5.635±0.2207) nuclei based on Perimeter values acquired from lamin B stained tissue images after pre-processing. **A. 1**.Comparative distribution of the number of normal (Mean±Std error of mean:23.47± 0.7129) (Std Dev:13.11±0.0541) and ovarian cancer (Mean±Std error of mean:51.62±1.153) (Std Dev:22.86±0.8153) nuclei based on Area values acquired from lamin A stained tissue images before pre-processing. **B. 2**. Comparative distribution of the number of normal (Mean±Std error of mean:19.94± 0.316) (Std Dev:8.414±0.2234)and diseased (Mean±Std error of mean:49.31± 1.234) (Std Dev:26.35±0.8725)nuclei based on Area values acquired from lamin B stained tissue images before pre-processing. **B. 3**.Comparative distribution of the number of normal (Mean±Std error of mean:14.66± 0.3791) (Std Dev:6.577±0.268) and ovarian cancer (Mean±Std error of mean:41.38±1.021) (Std Dev:19.24±0.722) nuclei based on Area values acquired from lamin A stained tissue images after pre-processing. **B. 4**.Comparative distribution of the number of normal (Mean±Std error of mean:13.02± 0.3677) (Std Dev:6.401±0.26) and ovarian cancer (Mean±Std error of mean:42.44±1.006) (Std Dev:18.13±0.7111) nuclei based on Area values acquired from lamin B stained tissue images after pre-processing.

Overall, these measurements confirmed prominent alteration in morphology in the cancer nuclei or in the nuclei approaching malignancy with respect to the normal nuclei and gave a gross idea regarding the direction of change. This experiment concluded that morphometric alteration in form of altered distribution of lamins in nuclei can potentially be used as signatures to classify cancer and normal nuclei or to study the progress towards malignancy and this feature being unaltered post pre-processing, can probably be a potential classifier for the deep learning model to distinguish between cancer and normal nuclei.

### Training a Deep Hybrid Learner

After pre-processed images were acquired, we had to split the data into a training set and validation set with a split ratio of 75:25. The training set was used to train the supervised binary classification model and the validation set was used for hyper-parameter tuning to make sure that the model was not over-fitting on the training set and remained generalized. For training a Deep Hybrid Learner, we have first trained the 21 Layered CNN which is used to extract features. We have trained it for 250 epochs and the model learning and loss curves obtained are shown in **Figure 5**. The training and validation learning and loss curve clearly showed that the model was neither over-fitting, nor under-fitting and thus it showed a very good fit. In the model learning curve, we used AUC Score as the metric for determining the fitness of the algorithm. We observed from the learning curve that the training and validation AUC Scores gradually increased with training iterations or epochs and the maximum training score after 250 epochs was obtained as 0.998 and the validation score was obtained as 0.997. The model loss curves for the training and validation set showed that the training and validation loss was gradually decreasing with increase in epoch, which was an indication that the model was learning gradually with more training iteration. Absence of any statistically significant variance between training and validation loss indicated absence of any over-fitting. These scores and extremely minimal model loss values along with graphical interpretations from the learning and loss curve supported the fact that the model was quite well generalized and expected to perform well on the test dataset.

**Figure 5:**
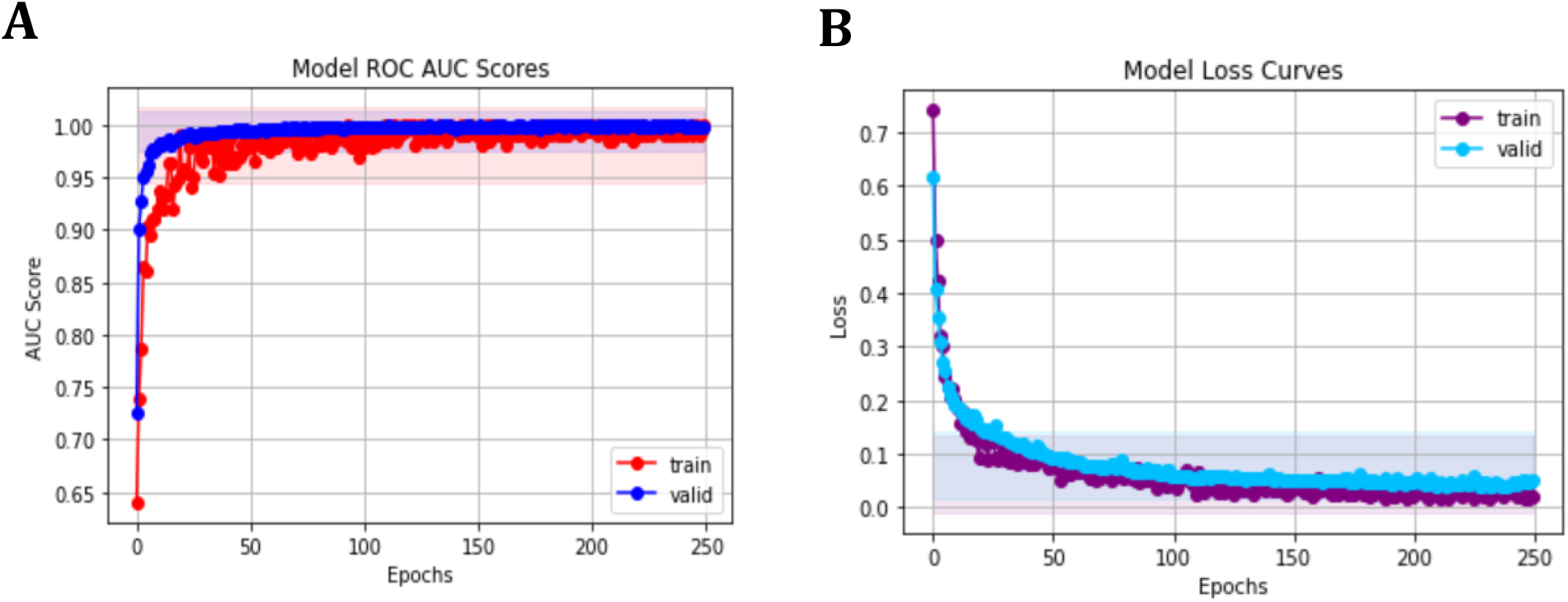
The plots detailing the pattern of. A. Model learning curve based on the information obtained from 250 epochs. Training AUC score has been depicted in red and validation AUC score has been depicted in blue. Both training and validation AUC scores have increased in every epoch .0.998 and 0.997 are the the maximum training and validation scores respectively after 250 epochs. **B. Model loss curve** based on the information obtained from 250 epochs. Training loss has been depicted in violet and validation loss has been depicted in blue. Training and validation losses are gradually decreasing in every epoch showing absence of any significant over-fitting.

### Selection of Model Evaluation Matrices

Since, we had a highly imbalanced dataset; accuracy alone would never be a good metric and could be misleading. If the model was always predicting the output as the majority class label, then the accuracy values would be very high, but yet the model would be highly biased on the majority class. Hence, we needed a metric that could show us the impact of true positives and false positives on an imbalanced dataset. Hence, we used Area under the Receiver Operating Characteristics Curve (AUC-ROC) scores (**Figure 5**) and Confusion matrix to clearly highlight the true positives, true negatives, false positives and false negatives, with the positive class being detection of cancer cells and the negative class being detection of normal cells. In our case, since AI driven approaches were used as automated pre-checks for cancer detection, after which detailed tests and inspection would be performed, false positives were comparatively less expensive than false negatives, as false negatives would lead to delayed detection of cancer. Typically then, a model with a specific set of hyper-parameters giving maximum AUC-ROC score, minimum false positives, and false negatives needs to be selected, in which less proportion of false negatives as compared to false positives will be preferred. .

### Model Evaluation on Test Data

The clinical details of the TMA samples were revealed and it contained a mixed cohort of tissues from Non cancer ovary as well as from omentum and adjacent areas from patients diagnosed with ovarian cancer undergoing debulking surgery. The normal and cancer tissue samples were grouped in 7 sub-classes (PDS adjacent Normal, PDS Tumor, IDS good response adjacent Normal, IDS good response Tumor, IDS poor response Normal, IDS poor response Tumor, Non cancer ovary). (**Figure 6)** One representative image from each sub-class was chosen as test image to evaluate the model performances. We compared performances of deep hybrid learning models (Deep Hybrid Learning with Random Forest(*45*) and Deep Hybrid Learning with XGBoost (*46*)) with other models like conventional deep neural network (DNN) model, Densenet 201(*47*) with transfer learning model(*48*), ResNet50(*49*) with transfer learning model(*48*), InceptionNetv3(*50*) with transfer learning model(*48*), VGG 16(*51*) with transfer learning model(*48*). The main difference between DHL and conventional DNN is that conventional DNN uses a fully connected neural network, whereas DHL uses a classical Machine Learning algorithm for final classification. DNN uses a fully connected neural network. In the transfer learning models, we have used pretrained weights from ImageNet(*52*). The training, validation and test dataset was the same for all the approaches and the number of epochs, batch size was also consistent for all the approaches.Confusion matrices were obtained for the 7 representative images from 7 sub-classes. The Deep Hybrid Learners exhibited 100% validation accuracy by recognising 4 images from 4 sub-classes (PDS adjacent Normal, IDS good response adjacent Normal, IDS poor response Normal and Non-Cancer Ovary, respectively) as true negatives or Normal and 3 images from 3 sub-classes (PDS Tumor, IDS good response Tumor, IDS poor response Tumor, respectively) as true positive or Cancer, whereas the other models could not recognise all 7 images accurately resulting in lower validation accuracies. ‘Normal’ and ‘Cancer’ has been annotated as 0 and 1 respectively in the confusion matrices. These images were absolutely unknown to the model and the clinical details were not revealed to the person performing the tests to ensure an unbiased validation and impartial selection of the accurate model based on performance. The matrices were used as the score cards to evaluate the model performances. So, deep hybrid learners were found to be the best working models for this specific problem. (**Figure 7**). For this research work, the choice of the ideal model architecture depends on two main criteria: Generalization and Efficiency. A model is said to be generalized when it is not over-fitting on training data and the model evaluation scores are consistent for training set, validation set and testing set. From the above results we can see that the Deep Hybrid Learners (both Random Forest and XGBoost variant) showed almost consistent results for training, validation and testing phases. Also, we found that the model was extremely efficient with low variance, as we observed that the AUC scores on training validation and test dataset were 0.99, 0.99 and 1.0 respectively (**Table 1**). The conventional deep learning model trained from scratch without transfer learning, seemed to have high training scores, but it showed high variance on validation and test dataset as the scores were much lower on validation and test set. Therefore, it indicated that the model was over fitting on the training data, and it was not generalized, hence performing poorly on testing and validation dataset. This behaviour of the model could be explained by our previous hypothesis that the dataset used for this research work was not favourable for a conventional deep learning approach, as it would require more training samples for the conventional model to learn and improve generalization. Hence, more sophisticated and novel approaches like Deep Hybrid Learning which uses CNN for feature extraction and classical machine learning algorithms for the final classification, was more efficient and robust for this type of microscopic image datasets. We have even applied Transfer Learning(*48*) with more sophisticated Deep Learning architectures like DenseNet201(*47*), ResNet50 (*49*), InceptionNetv3(*50*), VGG16(*51*), but the results obtained showed presence of over-fitting, lack of generalization and much lower model efficiency than the Deep Hybrid Learners. One plausible reason could be that these transfer learning models were trained using pre-trained weights from ImageNet (*52*) images, which were significantly different and might have a significant statistical difference from microscopic images, making transfer learning approach ineffective in this case. Thus we can conclude that our Deep Hybrid learning approach was successful and much better performing than other deep learning algorithms with these types of microscopic image datasets for automated detection of ovarian cancer cells.

**Figure 6:**
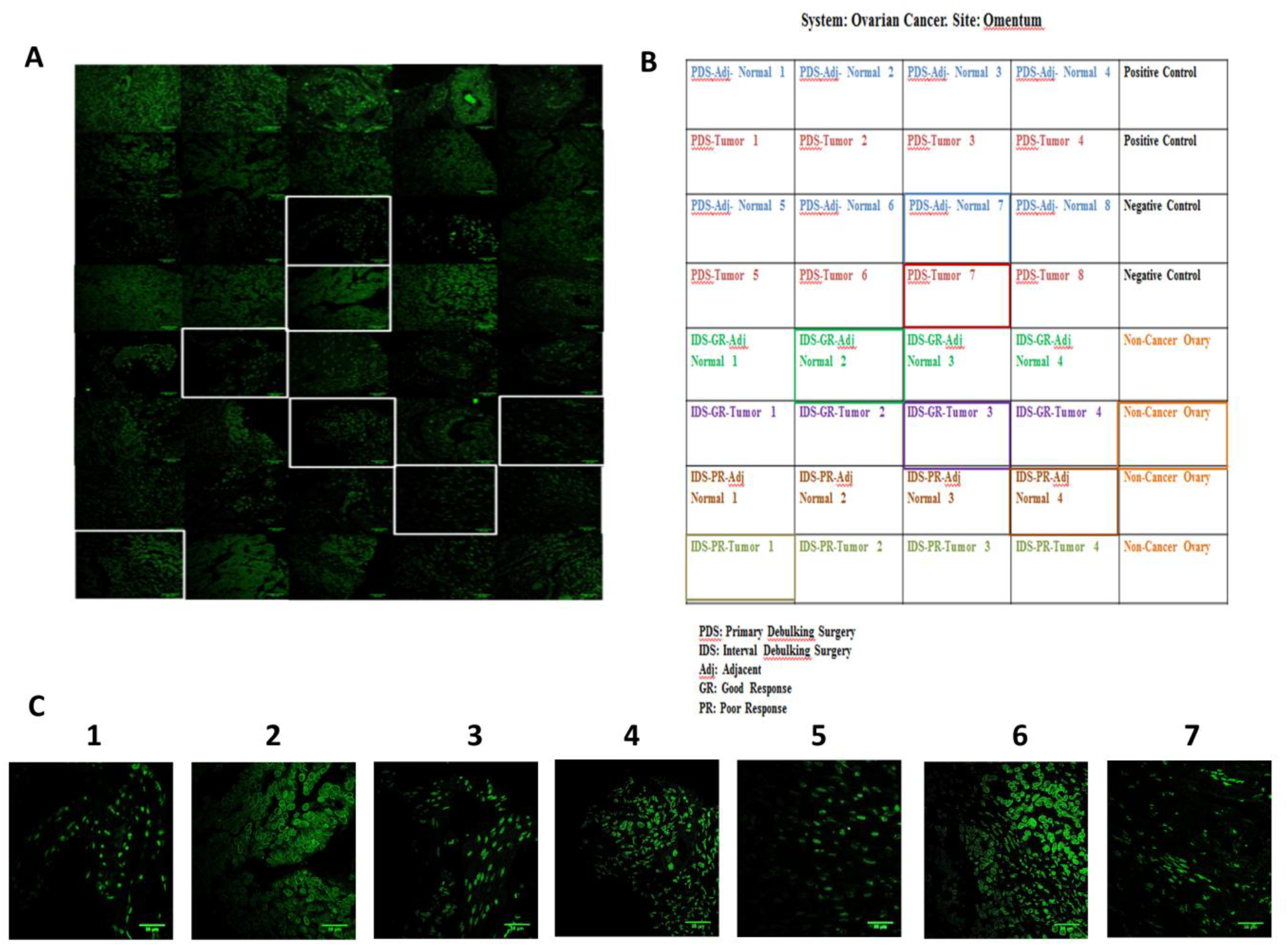
40 tissue samples from the TMA containing both normal and cancer tissues were grouped in 7 sub-classes and one representative image from each sub-class was used as test image for validation of model performances. **A**. 40 samples in a tissue microarray slide stained with lamin A. Representative images from 7 sub-classes have been marked in white boxes. Each field has been captured under Magnification: 63x. Scale Bar: 35 µm. **B**. Clinical details of microarray sample tissues from Non cancer ovary as well as from omentum and adjacent areas from patients diagnosed with ovarian cancer undergoing debulking surgery. 7 sub-classes have been marked in 7 different colours. Positive and Negative controls have been marked in black. C. Seven representative images which were used as test data for validation of the best working model. All normal and cancer tissues were grouped in seven sub-classes namely, 1. PDS-Adj-Normal 2.PDS-Tumor 3.IDS - GR-Adj Normal 4.IDS-GR-Tumor 5.IDS-PR-Adj Normal 6.IDS-PR-Tumor 7. Non-Cancer Ovary

**Figure 7:**
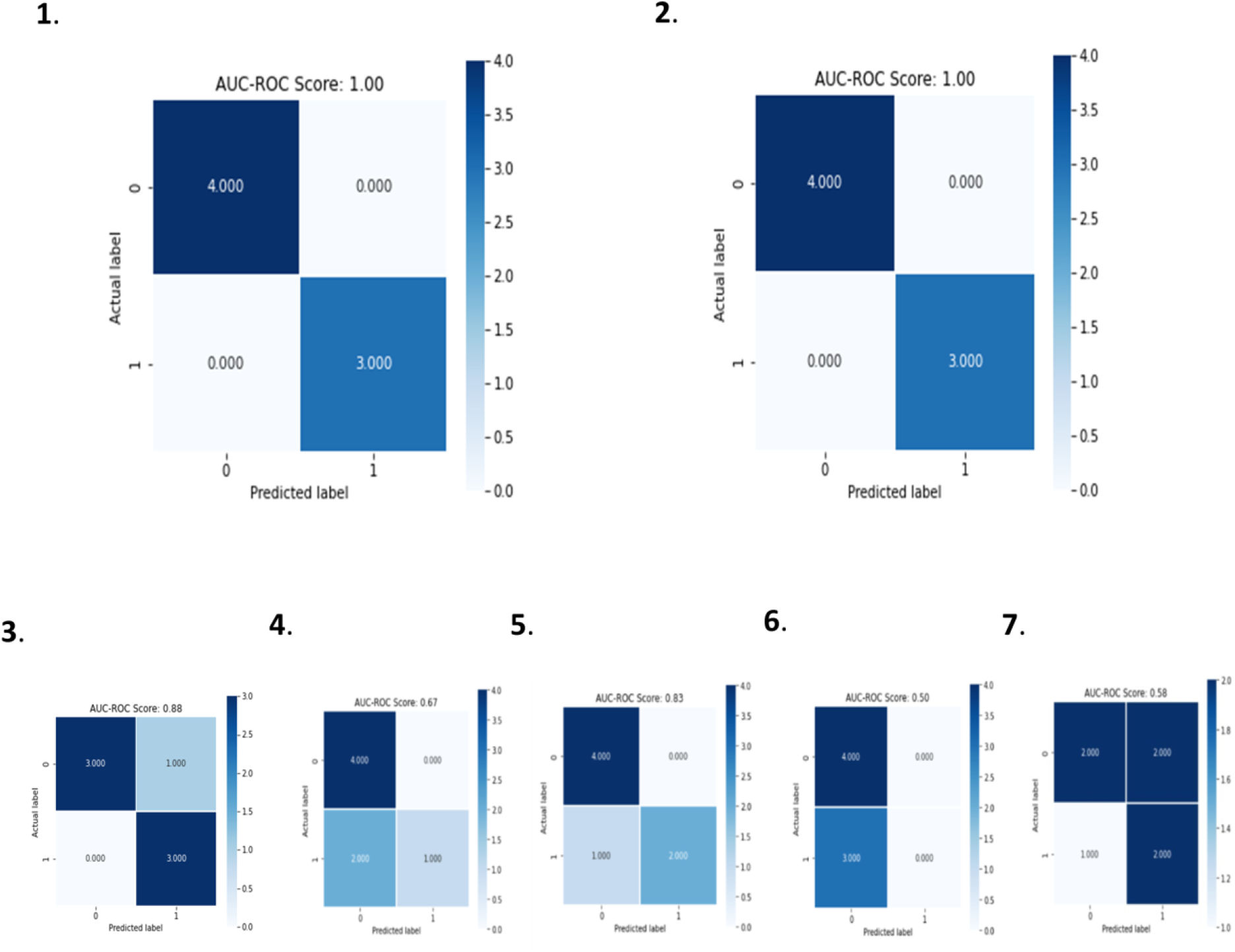
Confusion Matrix on Test Dataset. In this research work we have compared the Deep Hybrid Learning with both XGBoost and Random Forest variant, with a conventional Deep Neural Network (without transfer learning and having the same 21 layered CNN as DHL), DenseNet201 with transfer learning, InceptionNet v3 with transfer learning, ResNet50 with transfer learning and VGG16 with transfer learning. The Deep Hybrid Learners exhibited 100% validation accuracy by accurately idetifying 4 images from 4 classes (PDS adjacent Normal, IDS good response adjacent Normal, IDS poor response Normal and Non-Cancer Ovary, respectively) as true negatives or Normal and 3 images from 3 classes (PDS Tumor, IDS good response Tumor, IDS poor response Tumor, respectively) as true positive or Cancer. The other models could not identify all 7 images correctly resulting in lower validation accuracies. ‘Normal’ and ‘Cancer’ has been denoted as 0 and 1 respectively in the confusion matrices. **1**. Deep Hybrid Learning with Random Forest **2**. Deep Hybrid Learning with XGBoost**3**.Conventional Deep Neural Network Model **4**.DenseNet201 with transfer learning **5**.ResNet50 with transfer learning **6**.InceptionNetv3 with transfer learning **7**. VGG16 with transfer learning

### Deep Hybrid Learning

The trained and hyper-parameter tuned model performance on the test set proved how well the model was generalized and did not have any unwanted bias. Now, as imbalanced dataset is not suitable for classical deep learning models for building supervised classifiers with high accuracy and generalization, we came up with the Deep Hybrid Learning (DHL) algorithm, which utilized Deep Convolutional Neural Network to extract features from the pre-processed samples and uses the extracted feature vector with classical Machine Learning algorithms like Random Forest(*45*) and XGBoost (*46*) to build the final classifier. Of late Ensemble learning techniques like Boosting algorithms are known to work well with high dimensional data, as boosting techniques are known to combine weak learners to identify the “hard” data points and combine the weak learners to form a very strong and efficient classifier(*53*). Similarly, Ensemble methods like Random Forests work very well on smaller but high dimensional datasets for solving binary classification problems and have been known to produce generalized results(*45*). The results obtained using the Deep Hybrid Learning approaches turned out to be extremely promising (**Figure 7, Table 1**) and so far have performed much better than any other conventional approaches and is comparable to or even better than human level performance for this classification problem.

## Discussion

Application of convolutional neural network pipelines in cancer diagnosis and detection has already been widely recognised and reported by different groups utilising different technical and biological parameters. Currently, rise in life expectancy is a global concern which is widely being triggered by increase in incidences in age related gynaecological cancers (*54*). Morphometric studies in context of cancer has been thus far evaluated in only a few other carcinomas but not in a great detail in ovarian cancer. Immunocytochemical staining is a widely used method for detection and diagnosis of various cancers using different markers(*55*). On the other hand, alterations in nuclear morphometry have widely been acclaimed as hallmarks in various cancers and are used majorly in pathology or diagnosis purposes by clinicians to verify the degree of malignancy(*56-58*). But these verifications are potentially based on visual inspections and considering cellular heterogeneity in a tissue and as a result, an apparent subtlety in the changes post malignancy, manual observations are never trustworthy. To address this fallacy, recent researches have focussed on fluorescence imaging combined with machine learning for detection of malignancy. Various parametric and non-parametric methods are already in use for diagnosis of various cancers(*59-61*). Several groups have reported deep learning techniques to classify different stages or subtypes of ovarian cancer based on different techniques and biological features(*37, 39*). These rely on different classifiers based on differential features due to shift from the familiar genomic or proteomic sketch of the non-cancerous counterpart. There are several methods to study alterations in various nuclear matrix proteins which are already reported to be used for cancer diagnosis like BLCA4 (Bladder and urothelial carcinoma protein 4) in bladder cancer(*62*), PC1 (Prostate Cancer1) in prostate cancer(*63*) and NMP 179 (Nuclear Matrix Protein 179) in cervical cancer (*64*). But we have introduced nuclear lamins (lamin A and lamin B), which are residents of nuclear boundary and are ‘guardians’ of the nuclei controlling nuclear morphology. Hence their altered expression or distribution acts as a function of nuclear shape and size and has been used as the tool to detect and diagnose malignancy in context of ovarian cancer. Confocal images of tissue samples elucidated a significant increase in area and perimeter in the ovarian cancer nuclei which is in well agreement to the fact that the cell nuclei in the ovarian tumor tissues are mostly associated with an enlargement in size compared to the cell nuclei of normal tissues. Progressively, we increased the sample size and attempted to evaluate the possibility of quantitative feature extraction of nuclei and characterisation by projecting nuclear morphometry as a potential tool to distinguish normal and ovarian cancer tissues by introducing a novel deep hybrid learning network. We first focussed on extracting the pattern of the images and then moving a step further, noise reduction was performed so that it becomes convenient for the model to distinguish between cancer and normal tissue images. Pre-processing the images using various advanced mathematical and morphological techniques, like sharpening, masking, smoothening etc. enhanced the differentiating pattern of the images so that the neural model could easily identify them with utmost precision. This was followed by advanced techniques of data augmentation to create all sort of simulated practical tests by training the deep neural network model over the microscopic image dataset, using the Deep Hybrid learning approach, resulting in a much more reliable system than standard CNN. But, it’s a challenging problem to custom tailor the process of feature extraction in order to classify cancer from microscopic image dataset with fastness and precision. On the other hand, modern techniques of deep learning have already proven its superiority to perform image classification with minimal pre-processing and in many cases it has outperformed domain experts in classifying images. Thus, we decided to use Deep Learning approaches based on CNN (Convolutional neural network) for our study. A typical CNN involves one input and one output layer along with one or multiple hidden layers where convolution occurs through multiplication or other dot products and they are all connected with several nonlinear activation functions(*65*). For this specific problem, we believe that the distribution of lamins and size of the nucleus are important factors to classify images with cancer and normal nuclei. Thus, we have trained a deep learning network specifically crafted *de novo* for the particular challenge of classifying ovarian cancer.

Different types of augmentation bring almost all possible kinds of representation of the images resulting in a much higher possibility that a real time test image would be very similar if not exactly same and would be having greater probability to be classified with perfection in real time. Also, the drop out used here was 0.20 which is a regularization technique used for preventing over fitting of models. We have introduced a combination of classical machine learning algorithms (XGBoost, SVM, Random Forest) with standard CNN and designed a state of the art deep hybrid network. We created a completely automated pipeline architecture starting from processing images to predicting a cancer or a non-cancer which would be performed within seconds. The model showed 99.8% training accuracy and 99.7% validation accuracy in distinguishing normal and ovarian cancer cell nuclei and with our feedback mechanism the network could be retrained with wrong predictions made to further improve the accuracy. So a more robust model could be made which outperformed clinical precision in terms of speed and accuracy. As the problem is as unique as ovarian cancer detection using structures / patterns of images, we did not rely over the pre-trained CNN models rather we focused to train a deep learning model tailor-made for solving this particular problem and made it much more operational in real life circumstances rather restricting it to be an academic research tool. As part of our experiment, we used transfer learning models like ResNet50, AlexNet, VGG-16, DenseNet and our custom model had outperformed all of them. That is why it was obvious that the transformation and feature engineering was the main differentiator for solving this particular problem. The model has worked perfectly well and it is certain that with sufficient images in future this would outperform other models and will be able to classify not only between normal and cancer nuclei but will also be able to predict the degree of risks of benign nuclei to become cancerous.

Most female genital tract malignancies possess identifiable precursors except for ovarian malignancies which is the 8th leading cause of cancer mortality among women and 7th leading cause of cancer diagnosis worldwide(*66*). In this alarming scenario, an early detection method for ovarian cancer is significantly pressing, as ovarian carcinoma currently do not have a screening modality, also the symptoms are not very unique and only arise after the disease is in advanced stage (*54*).We know that majority of HGS (High grade serous) ovarian cancers arise from fallopian tubes. Many women undergo fimbriectomy for sterilisation purposes as well as a prophylactic measure for prevention of ovarian cancer in high risk individuals i.e., BRCA mutations. It would be interesting to study using the same method in the future whether alteration in the nuclear architecture in fimbria / ovary could be one of the early predictors for developing cancer. More importantly, this may be used as prognostic marker in addition to the standard immunohistochemistry and histology in ovarian cancer if a clinical correlation can be shown in future studies and that remains one of our target research strategies in the future including application in other women cancers. A lot of studies are trying to detect precursor lesion signatures like STIC (serous tubal intraepithelial cancer and p53 signature in ovarian/tubal cancer; NUMODRIL approach can add to these models for improving diagnostic accuracy. Further application in ovarian cancer could be supplementing chemotherapy response score (CRS) after neoadjuvant chemotherapy and IDS, where CRS 2 score is the grey zone and requires better biomarkers for prognostic stratification.

The novelty of this work lies in the fact that this pipeline christened as NUMODRIL could be generalized as a global technique in the diagnosis and prognosis of all types of neoplasia which might be associated with changes in nuclear architecture as a result of alteration of lamin A/B expression profile.

## Materials and Methods

### Tissue sample collection

Formalin fixed paraffin embedded tissues from ovarian cancer patients (n=40) were obtained during frontline surgery at Tata Medical Centre (TMC), Kolkata. Ethical approval for the study was obtained (EC/TMC/45/15). An ovarian tissue micro array was created at TMC representing various stages of high grade serous ovarian cancer and consisting of both Primary debulking surgery and interval debulking surgery tissues. Slides were provided without identifying or any clinical data for unbiased analysis.

### Immunohistochemistry

All the tissues of normal ovary and ovarian cancer and Tissue microarray samples were obtained from Tata Medical Centre following the ethical guidelines. The paraffin embedded blocks were cut into 4-5 µm sections and affixed onto glass slides. For removal of paraffin, the slides were immersed in xylene (3*10mins) followed by immersion in graded ethanol twice for 10 minutes in each (100%,95%,80%,70%), washed in ddH_2_O twice 5 minutes each. The tissues were immersed in 10mM sodium citrate buffer (pH 6), placed in a microwavable container, and heated at full power for 3 minutes and 80% power for subsequent 12 minutes. After which the slide was allowed to gradually cool in the same buffer, followed by rinsing with ddH_2_O twice for 5 minutes each followed by two rounds of wash in PBS for 5 minutes each. Tissues were permeabilised with 0.5% Triton X 100 for 5 minutes, blocked and incubated with primary antibody solution containing blocking agent (5% Normal Goat Serum and Primary antibody as per dilution recommended in 1X PBS) in a humidified environment for 2 hours in room temperature. Subsequently cells were incubated with secondary antibody diluted in 1X PBS for 2 hours in dark in room temperature. Cells were finally incubated in Propidium Iodide at a concentration of 10μg/ml for 40 minutes and then mounted on glass slides with mounting medium containing anti oxidizing agent (PPD). It was sealed with nail polish and imaged using 63X oil immersion objectives in Zeiss LSM 710 Meta confocal Microscope. The slides were stored in dark at 4°C for further use. Primary antibody dilutions for Rabbit Anti Lamin A antibody (Sigma Aldrich L1293), Anti Lamin B (Santacruz sc-6217) Antibody were 1:100 and 1:50 respectively. Secondary antibodies were conjugated with Alexa Fluor 488 (Green Fluorescence) and used at a dilution of 1:400. For Anti Lamin B antibody, methanol fixation was performed. Cells were fixed with ice cold methanol for 20 minutes at −20°C and permeabilised in Acetone for 1 minute in room temperature.

### Image Analysis and data presentation

Images were analysed using ImageJ software (ImageJ bundled with 64-bit Java 1.8.0_112). Considering each nucleus an ellipse, the equations used in the referred article (*44*) were followed to derive the values of the morphometric parameters like area, perimeter, loop length, circularity, eccentricity, foci distance, maximum curvature, normalized maximum curvature. Histograms were generated using ROOT: An object oriented C++ framework for large data storage, presentation, visualisation and statistical analysis (Version 6, Release 6.08/06-2017-03-02). Each field from every tissue sample contained around 100 nuclei approximately. The length of the major and minor axes of each of the nuclei (around 1000 normal nuclei and 860 diseased nuclei) was measured manually by ImageJ. Considering each nuclei an ellipse, the morphometric parameters like area, perimeter, loop length, circularity, eccentricity, foci distance, maximum curvature, normalized maximum curvature were calculated. Histograms were generated using ROOT: An object oriented C++ framework for large data storage, presentation, visualisation and statistical analysis (Version 6, Release 6.08/06-2017-03-02).). In the plots the X axis denotes the normalized number of nuclei with respect to the total number of nuclei calculated corresponding to the defined parameter and Y axis denotes the measure of the parameter. Mean, Standard error of mean and Standard deviation for the analysis of each parameter have been mentioned in the figure legends.

### Pre-processing

Although Deep Learning algorithms are believed to apply auto feature extraction methods, we wanted to additionally transform the raw data and make it easier for the model to unravel key features. As illustrated in **Figure 3** using Morphological elliptical Image filters, morphological masks were formed around the nuclei and then using Gaussian Blur filter and pixel weights addition, the sharpened version of the images were obtained. Then the images were grey-scaled and normalized before feeding it into the deep learning model.

### Synthetic Minority Oversampling Technique (SMOTE)

Synthetic Minority Oversampling Technique (SMOTE)(*40*), uses vector interpolation techniques with high dimensional data to generate synthetic samples of the minority class. The synthetically generated samples are linear combinations of two similar samples from the minority class (x and x^R^) and are defined as:

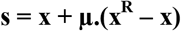

where, **x** is a minority class sample and **x**^**R**^ is another minority class sample which is randomly picked from a sample space which has high similarity with **x** and is identified by nearest neighbour approach. The factor **µ** is randomly selected from a uniform distribution between 0 and 1 and it is the same for all dimensional variables, but it is different for different values of x. SMOTE keeps the same expected value of the synthetically generated minority class, but it decreases its variability, thereby making the SMOTE-augmented data statistically different and thus, does not add any bias to the data.

### Deep Hybrid Learning Model Architecture

The entire Deep Hybrid Learning architecture used for this research work starts with the pre-processing layer, then the pre-processed images are passed through the model input layer. Each input is of dimension (1024, 1024,4). Deep Hybrid Learner uses a Deep Convolutional Neural Network Layer for feature extraction. In our research we have used a 21 Layered CNN which is inspired from InceptionNet v3(*50*) architecture. The CNN part consists of a series of Incept layer and Squeeze Layer which are like grouped Convolution Layers with specific hyper-parameters and the nested Conv2D layers is illustrated in **Supplementary figure 2**.

For the Incept Layer, it takes the number of filters for the Conv2D sub-layer and another hyper-parameter for the number of filters for Left and Right Conv2D sub-layer as input. In both the Conv2D sub-layers, after tuning, we have used a learning rate of 0.1 and an activation function of Leaky ReLu to learn the non-linear relationships in the underlying high dimensional data. For the initial Conv2D layer, we have used a filter dimension of 5×5 and used strided convolution with stride as 2. For Left Conv2D a filter size of 3×3 is used and for the Right Conv2D a filter size of 5×5 is used. Finally, both the left and right conv2d sub layers are concatenated and passed to the next layer. The Squeeze layer follows the same structure as Incept Layer. The learning rates and the activation function used in the sub-layers is the same, but the only difference is with the filter dimensions. For the initial Conv2D sub layer, the dimension is (1×1) and stride 1. While for Left Conv2D the filter dimension is 1×1 and for Right Conv2D the filter dimension is 3×3. Like the Incept layer, the Left and the Right Sub-layers are concatenated and passed to the next layers. After a series of Incept and Squeeze layers we have used another Conv2D layer with 64 filters and each filter of dimension 3×3, with an activation function of Leaky ReLu with learning rate 0.1. Finally, after all the convolution layers which are used to extract the features, we have flattened the output and passed the flattened output to classical Machine Learning algorithms like XGBoost and Random Forest for the final classification part. The overall scheme of deep hybrid learning used here is depicted in **Supplementary figure 2**.

### Comparison of DHL with other Deep Learning approaches

In order to evaluate how well our Deep Hybrid Learning approach is performing on the training, validation and test dataset as compared to other popular Deep Learning approaches, we have used AUC Scores, Accuracy and Confusion Matrix as the evaluating metrics. In this research work we have compared the Deep Hybrid Learning with both XGBoost and Random Forest variant, with a conventional Deep Neural Network (without transfer learning and having the same 21 layered CNN as DHL), DenseNet201(*47*) with transfer learning, InceptionNet v3(*50*) with transfer learning, ResNet50(*49*) with transfer learning and VGG16(*51*) with transfer learning.

## Acknowledgements

All tissues and TMA samples were obtained from Tata Medical Centre, provided by AsimaMukhopadhyay, a clinician scientist (Gynaecological Oncologist), based at Tata Medical Centre and currently at ChittaranjanNational Cancer Institute Kolkata and her research group. Biobanking for ovarian cancer tissues and TMA creation was performed through the DST-UKIERI grant held by Dr.AsimaMukhopadhyay. The histograms were generated using ROOT data analysis framework with the help of Mr.Gourab Saha, SRF, High Energy Nuclear and Particle Physics Division, SINP who helped in writing the programs in C++. Development of CNN and the following analysis based on that was performed by Sk. Nishan Ali and Aditya Bhattacharya, MUST Research. Duhita Sengupta thanks DAE for the fellowship. Kaushik Sengupta thanks SERB, DST& BARD project of DAE, Govt. of India.

## Authors Contributions

Tissue samples were provided by Asima Mukhopadhyay as a part of the collaborative study HGSC Lamin with Kaushik Sengupta. Duhita Sengupta performed the major experiments. SkNishan Ali, Aditya Bhattacharya and Joy Mustafi developed the CNN. Duhita Sengupta, Kaushik Sengupta, Sk Nishan Ali and Aditya Bhattacharya wrote the manuscript. KaushikSengupta conceived the entire project. Asima Mukhopadhyay reviewed the manuscript and provided clinical context to application in ovarian cancer.

## Supplementary Figures and Tables

**Figure 1:**
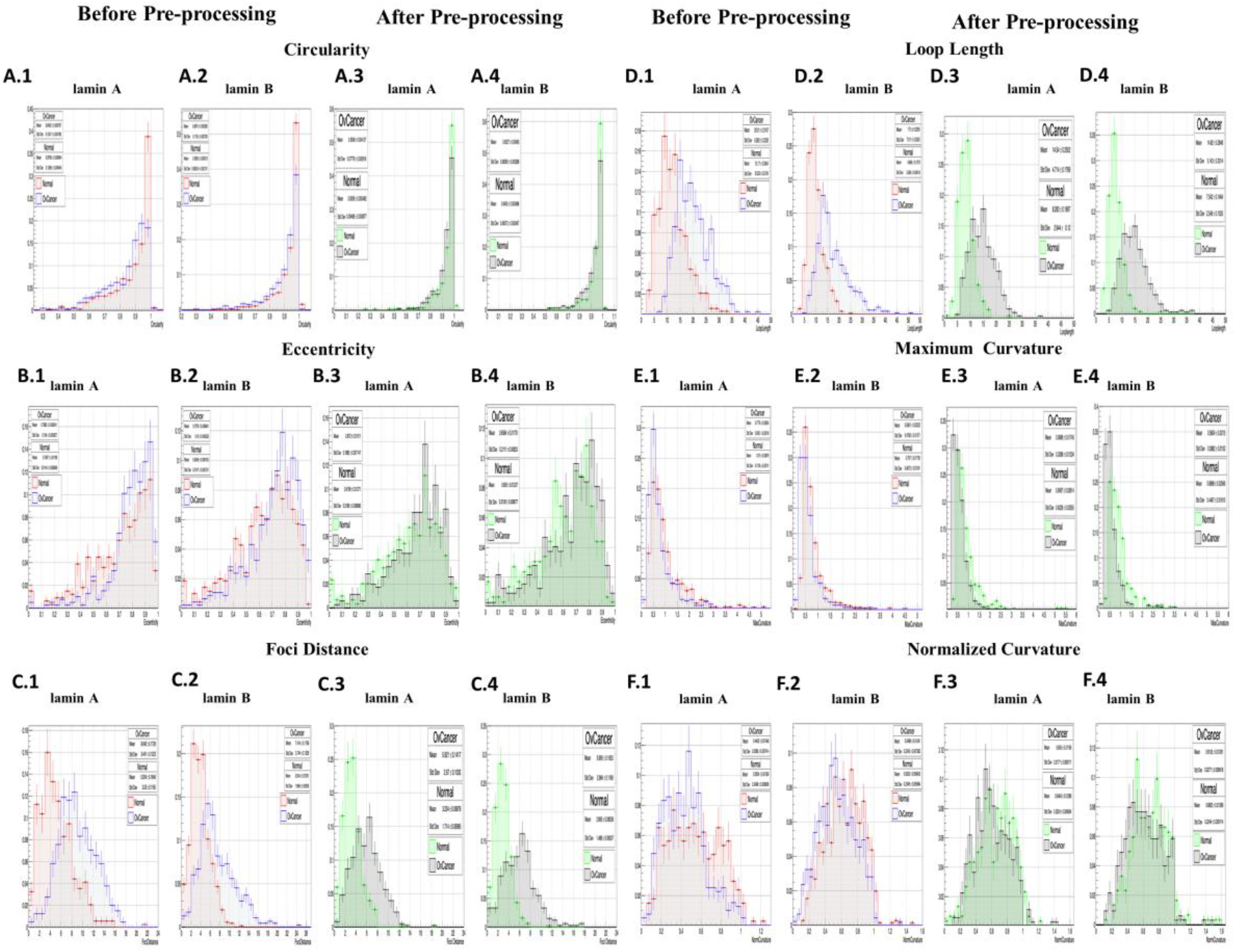
Histograms showing distributions of the normal and Ovarian Cancer nuclei based on different morphometric parameters obtained from lamin A and B stained tissue sample images before and after pre-processing. X axis denotes the normalised number of nuclei with respect to the total number of nuclei calculated. Y axis denotes the measure of the parameter. **A. 1**.Comparative distribution of the number of normal (Mean±Std error of mean:0.8796± 0.005994) (Std Dev:0.1285±0.00495) and ovarian cancer (Mean±Std error of mean:0.8452± 0.006767) (Std Dev:0.1347±0.004785) nuclei based on Circularity values acquired from lamin A stained tissue images before pre-processing. **A. 2**. Comparative distribution of the number of normal (Mean±Std error of mean:0.9309± 0.003013) (Std Dev:0.8024±0.002131) and ovarian cancer (Mean±Std error of mean:0.8974± 0.005269) (Std Dev:0.1135±0.003726) nuclei based on Circularity values acquired from lamin B stained tissue images before pre-processing. **A. 3**. Comparative distribution of the number of normal (Mean±Std error of mean:0.9309± 0.005482) (Std Dev:0.09496±0.003877) and ovarian cancer (Mean±Std error of mean:0.9248± 0.004127) (Std Dev:0.0776±0.002918) nuclei based on Circularity values acquired from lamin A stained tissue images after pre-processing. **A. 4**. Comparative distribution of the number of normal (Mean±Std error of mean:0.9455± 0.002488) (Std Dev:0.06072±0.002467) and ovarian cancer (Mean±Std error of mean:0.9227± 0.004665) (Std Dev:0.08395±0.03328) nuclei based on Circularity values acquired from lamin B stained tissue images after pre-processing. **B. 1**.Comparative distribution of the number of normal (Mean±Std error of mean:0.7067± 0.01169) (Std Dev:0.2144±0.008269) and ovarian cancer (Mean±Std error of mean:0.7885± 0.008241) (Std Dev:0.164±0.005827) nuclei based on Eccentricity values acquired from lamin A stained tissue images before pre-processing. **B. 2**. Comparative distribution of the number of normal (Mean±Std error of mean:0.6348± 0.008105) (Std Dev:0.2147±0.005731)and diseased (Mean±Std error of mean:0.7076± 0.008841) (Std Dev:0.19±0.006252)nuclei based on Eccentricity values acquired from lamin B stained tissue images before pre-processing. **B. 3**.Comparative distribution of the number of normal (Mean±Std error of mean:0.6196± 0.01271) (Std Dev:0.2186±0.00896) and ovarian cancer (Mean±Std error of mean:0.672±0.01011) (Std Dev:0.1895±0.0071471) nuclei based on Eccentricity values acquired from lamin A stained tissue images after pre-processing. **B. 4**.Comparative distribution of the number of normal (Mean±Std error of mean:0.605± 0.01227) (Std Dev:0.2108±0.00867) and ovarian cancer (Mean±Std error of mean:0.6584±0.01178) (Std Dev:0.2111±0.00833) nuclei based on Eccentricity values acquired from lamin B stained tissue images after pre-processing. **C. 1**.Comparative distribution of the number of normal (Mean±Std error of mean:5.204± 0.1648) (Std Dev:3.03±0.1165) and ovarian cancer (Mean±Std error of mean:8.942± 0.1729) (Std Dev:3.441±0.1223) nuclei based on Foci Distance values acquired from lamin A stained tissue images before pre-processing. **C. 2**. Comparative distribution of the number of normal (Mean±Std error of mean:3.934± 0.07391) (Std Dev:1.968±0.05226) and diseased (Mean±Std error of mean:7.414± 0.1738) (Std Dev:3.744±0.1229)nuclei based on Foci Distance values acquired from lamin B stained tissue images before pre-processing. **C. 3**.Comparative distribution of the number of normal (Mean±Std error of mean:3.234± 0.09879) (Std Dev:1.714±0.06985) and ovarian cancer (Mean±Std error of mean:5.921±0.1417) (Std Dev:2.67±0.1002) nuclei based on Foci Distance values acquired from lamin A stained tissue images after pre-processing. **C. 4**.Comparative distribution of the number of normal (Mean±Std error of mean:2.895± 0.8538) (Std Dev:1.486±0.06037) and ovarian cancer (Mean±Std error of mean:5.995±0.1653) (Std Dev:2.984±0.1169) nuclei based on Foci Distance values acquired from lamin B stained tissue images after pre-processing. **D. 1**.Comparative distribution of the number of normal (Mean±Std error of mean:12.17± 0.3004) (Std Dev:5.523±0.2124) and ovarian cancer (Mean±Std error of mean:20.01± 0.3152) (Std Dev:6.282±0.2232) nuclei based on Loop Length values acquired from lamin A stained tissue images before pre-processing. **D. 2**. Comparative distribution of the number of normal (Mean±Std error of mean:9.869± 0.1275) (Std Dev:3.396±0.09018)and diseased (Mean±Std error of mean:17.5± 0.3255) (Std Dev:7.011±0.2302)nuclei based on Loop Length values acquired from lamin B stained tissue images before pre-processing. **D. 3**.Comparative distribution of the number of normal (Mean±Std error of mean:8.263± 0.1697) (Std Dev:2.944±0.12) and ovarian cancer (Mean±Std error of mean:14.54±0.2502) (Std Dev:4.714±0.1769) nuclei based on Loop Length values acquired from lamin A stained tissue images after pre-processing. **D. 4**.Comparative distribution of the number of normal (Mean±Std error of mean:7.542± 0.1464) (Std Dev:2.549±0.1035) and ovarian cancer (Mean±Std error of mean:14.82±0.2848) (Std Dev:5.143±0.2014) nuclei based on Loop Length values acquired from lamin B stained tissue images after pre-processing. **E. 1**.Comparative distribution of the number of normal (Mean±Std error of mean:1.1015± 0.03876) (Std Dev:0.7105±0.02741) and ovarian cancer (Mean±Std error of mean:0.7776± 0.02854) (Std Dev:0.5651±0.02018) nuclei based on Maximum Curvature values acquired from lamin A stained tissue images before pre-processing. **E. 2**. Comparative distribution of the number of normal (Mean±Std error of mean:0.797± 0.01755) (Std Dev:0.4672±0.01241)and diseased (Mean±Std error of mean:0.6361± 0.0222) (Std Dev:0.4782±0.01571)nuclei based on Maximum Curvature values acquired from lamin B stained tissue images before pre-processing. **E. 3**.Comparative distribution of the number of normal (Mean±Std error of mean:0.9067± 0.03614) (Std Dev:0.6228±0.02555) and ovarian cancer (Mean±Std error of mean:0.5868±0.01745) (Std Dev:0.3288±0.01234) nuclei based on Maximum Curvature values acquired from lamin A stained tissue images after pre-processing. **E. 4**.Comparative distribution of the number of normal (Mean±Std error of mean:0.8898± 0.02566) (Std Dev:0.4467±0.01815) and ovarian cancer (Mean±Std error of mean:0.5804±0.0215) (Std Dev:0.3882±0.0152) nuclei based on Maximum Curvature values acquired from lamin B stained tissue images after pre-processing. **F. 1**.Comparative distribution of the number of normal (Mean±Std error of mean:0.5354± 0.01359) (Std Dev:0.2496±0.009609) and ovarian cancer (Mean±Std error of mean:0.4402± 0.01048) (Std Dev:0.2086±0.007414nuclei based on Normalized Curvature values acquired from lamin A stained tissue images before pre-processing. **F. 2**. Comparative distribution of the number of normal (Mean±Std error of mean:0.6323± 0.008435) (Std Dev:0.2246±0.005964)and diseased (Mean±Std error of mean:0.5489± 0.01401) (Std Dev:0.2243±0.007362) nuclei based on Normalized Curvature values acquired from lamin B stained tissue images before pre-processing. **F. 3**.Comparative distribution of the number of normal (Mean±Std error of mean:0.6464± 0.01286) (Std Dev:0.2224±0.009094) and ovarian cancer (Mean±Std error of mean:0.605±0.01156) (Std Dev:0.2177±0.008171) nuclei based on Normalized Curvature values acquired from lamin A stained tissue images after pre-processing. **F. 4**.Comparative distribution of the number of normal (Mean±Std error of mean:0.6803± 0.01289) (Std Dev:0.2244±0.009114) and ovarian cancer (Mean±Std error of mean:0.6103±0.01261) (Std Dev:0.2277±0.0.008918) nuclei based on Normalized Curvature values acquired from lamin B stained tissue images after pre-processing.

**Figure 2a:**
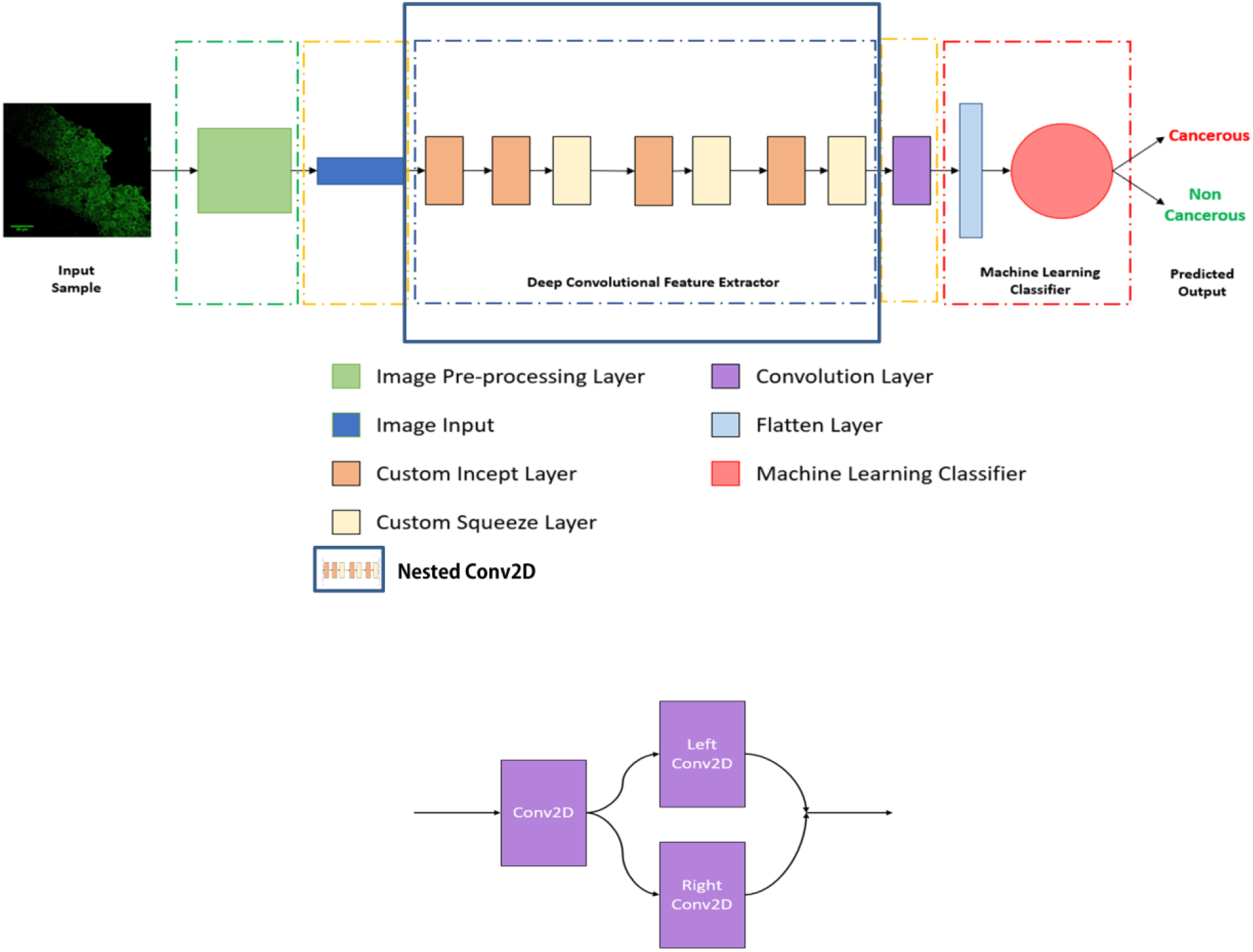
The overall scheme of deep hybrid learning algorithm.

**Table 1:** **Implication of different properties of SMOTE over high dimensional data**

| Property | Implications of SMOTE over high-dimensional data |
| --- | --- |
| $E(\text{SMOTE}) = E(X)$ | Minimal impact over the classifiers which depend upon mean values; |
| $\text{var}(\text{SMOTE}) = 2/3 * \text{var}(X)$ | Minority class variability is underestimated. Negatively impacts over classifiers which uses class-specific variances |
| $d(\text{SMOTE}, \text{TEST}) < d(X, \text{TEST})$: Euclidean distance | Test samples are classified mostly in the minority class for classifiers based on Euclidean distance. (variable selection can be handy in reducing this problem) |
| $\text{cor}(\text{SMOTE}, X) \geq 0$; $\text{cor}(\text{SMOTE}_s, \text{SMOTE}_t) \geq 0$ | Results in creating correlated samples thus, individual samples are not independent any further. |

